# MAPK/ERK signaling blocks ectopic H3K9me3 heterochromatin formation to confer mesoderm and endoderm developmental competence

**DOI:** 10.1101/2025.08.25.672203

**Authors:** Satoshi Matsui, Marissa Granitto, Samuel Sampson, Gerardo Mirizio, Ryo Maeda, Makoto Tachibana, Christopher Ahn, Hee-Woong Lim, Makiko Iwafuchi

**Author notes:** Present address: Premium Research Institute for Human Metaverse Medicine (PRIMe), The University of Osaka, Osaka 565-0871, Japan (Satoshi Matsui).

## Abstract

During gastrulation, dynamic interplay among cell signaling pathways dictates cell fate decisions. While extensive studies have elucidated their critical roles in morphological regulation, how these signals orchestrate the epigenome to confer developmental competence remains unclear. In this study, we demonstrate that H3K9me3-marked facultative heterochromatin domains undergo global reorganization during differentiation of human pluripotent stem cells into mesoderm and endoderm, which arise through epithelial-mesenchymal transition (EMT), but not into ectoderm, which retains epithelial state. We identify the MAPK/ERK pathway, acting downstream of FGF signaling, as a key mediator of this reorganization. Specifically, the MAPK/ERK pathway prevents ectopic formation of H3K9me3 domains at EMT– and lineage-specific gene loci whose expression is necessary for mesoderm and endoderm differentiation. Collectively, our findings reveal a previously unrecognized role for MAPK/ERK signaling in reorganizing the H3K9me3 landscape to enable mesoderm and endoderm differentiation, bridging a critical gap in our knowledge of how cell signaling pathways shape the epigenetic landscape during development.

## Introduction

During early embryonic development, epithelial epiblast cells differentiate into the three definitive germ layers – ectoderm, mesoderm, and endoderm – establishing the foundation of the body plan (1, 2). While the early-stage ectoderm retains its epithelial state, the endoderm and mesoderm undergo dramatic morphological changes, characterized by epithelial-to-mesenchymal transition (EMT) during gastrulation (3–6). Gastrulation is tightly regulated by BMP, WNT, NODAL, and FGF signaling pathways (7, 8). BMP4 signaling activates the WNT3 pathway, which in turn activates the NODAL pathway (9–11). Together with WNT3 and NODAL signaling, the FGF-mediated MAPK/ERK pathway plays a pivotal role in promoting EMT, cell migration, and the specification of mesoderm and endoderm (12–19). While FGF activity is essential for NODAL and WNT signal-mediated gene induction during gastrulation (13, 17, 20), the functional mechanisms of FGF/MAPK/ERK remain the least understood amongst the key gastrulation signals.

In addition to cell signaling, chromatin organization into heterochromatin and euchromatin through epigenetic mechanisms profoundly influences gene regulation, conferring transcriptional and developmental competency. Heterochromatin domains are less accessible to transcription factors and transcriptional machinery than euchromatin, acting as barriers to gene activation (21–23). These domains are predominantly marked by H3K9me3 and Polycomb repressive modifications, such as H3K27me3 and H2AK119ub1 (24–26). Polycomb repressive domains dynamically relocate to regulate key developmental genes, including *Hox* genes, to achieve proper embryogenesis (27–31). Although H3K9me3 has been traditionally associated with constitutive heterochromatin that stably represses repeat-rich genomic regions, emerging evidence reveals its dynamic role as a mark of facultative heterochromatin during embryonic development, where it contributes to lineage-specific gene regulation (26, 32, 33). For instance, during definitive endoderm differentiation into hepatic and pancreatic lineages in mouse embryos, H3K9me3-marked domains undergo dynamic remodeling, including both gains and losses at protein-coding loci (34). During early development, these facultative heterochromatin domains repress genes associated with mature cell functions, while their removal at later time points facilitates hepatic and pancreatic differentiation. Similarly, during *C. elegans* development, H3K9me3 is removed from differentiated cell-associated genes and gained at genes expressed earlier in development or in alternative lineages (35).

During mouse pre-implantation stages, *de novo* formation of H3K9me3 domains in the extraembryonic ectoderm antagonizes transcription factor binding to key developmental genes, serving as an effective repressive mechanism for embryonic lineages (36). Despite these functional significances of H3K9me3 facultative heterochromatin, the mechanisms driving its lineage-specific reorganization during cell differentiation remain poorly understood. In particular, how cell signaling pathways contribute to this dynamic heterochromatin reorganization during development has not been fully explored. One study reported that prolonged overactivation of the Raf/ERK signaling pathway in mouse fibroblasts altered the nuclear distribution of H3K9me3, as observed by immunofluorescent imaging (37). However, how physiological signaling impacts H3K9me3, especially its genome-wide localization during cell differentiation, remains to be elucidated.

To address these open questions, we utilized *in vitro* human germ layer differentiation models derived from hPSCs, directing differentiation toward paraxial mesoderm (PM), definitive endoderm (DE), and neural ectoderm (NE). We uncovered a notably more profound genome-wide reorganization of H3K9me3 heterochromatin domains during hPSC differentiation into PM and DE lineages that undergo EMT, compared to NE that remains epithelial. Furthermore, we applied a MEK inhibitor (MEKi) to suppress the MAPK/ERK signaling pathway during germ layer differentiation and found that the MEKi application impaired H3K9me3 domain reorganization and simultaneously compromised PM and DE differentiation. Specifically, in the absence of MAPK/ERK activity, ectopic formation of H3K9me3 domains occurred at EMT– and lineage-specific gene loci critical for mesoderm and endoderm differentiation. These findings highlight the crucial role of the MAPK/ERK pathway in reorganizing the H3K9me3 landscape to enable the induction of essential EMT– and lineage-specific genes for mesoderm and endoderm differentiation.

## Materials and Methods

### Reagents

Antibody information for immunostaining experiments.

| Antigen | Supplier | Cat no. | Lot no. | Host | Dilution factor |
| --- | --- | --- | --- | --- | --- |
| OCT4 | DSHB | PCRP-POU5F1-1D2 | N/A | Mouse | 1:20 |
| SOX2 | R&D Systems | AF2018 | KOY0419101 | Goat | 1:1000 |
| SOX2 | Cell Signaling | 4900S | 5 | Mouse | 1:500 |
| SOX1 | Cell Signaling | 4194S | 2 | Rabbit | 1:1000<br>1:250 |
| TBX6 | R&D Systems | AF4744 | CAPT0217111 | Goat | 1:200 |
| CDX2 | Cell Marque | 235R-14 | 0000027991 | Rabbit | 1:500 |
| FOXA2 | Abnova | H00003170-M1 | H0000317 | Mouse | 1:1000 |
| SOX17 | R&D Systems | AF1924 | KGA0818071 | Goat | 1:1000 |
| CDH1 | Cell Signaling | 3195S | 13 | Rabbit | 1:200 |
| CDH2 | Cell Signaling | 13116S | 4 | Rabbit | 1:200 |
| LAMINB1 | Abcam | ab16048 | GR3283674-1 | Rabbit | 1:1000 |
| anti-Mouse<br>Alexa488 | Jackson<br>ImmunoResearch | 711-545-152 | 135209 | Donkey | 1:500 |
| anti-Rabbit<br>Alexa488 | Jackson<br>ImmunoResearch | 711-545-152 | 138271 | Donkey | 1:500 |
| anti-Rabbit<br>Alexa594 | Jackson<br>ImmunoResearch | 711-585-152 | 136429 | Donkey | 1:500 |
| anti-Rabbit<br>Alexa647 | Jackson<br>ImmunoResearch | 711-605-152 | 138314 | Donkey | 1:500 |
| anti-Goat<br>Alexa594 | Jackson<br>ImmunoResearch | 705-585-147 | 130926 | Donkey | 1:500 |
| Phalloidin<br>Alexa647 | Abcam | ab176759 | N/A | - | 1:500 |

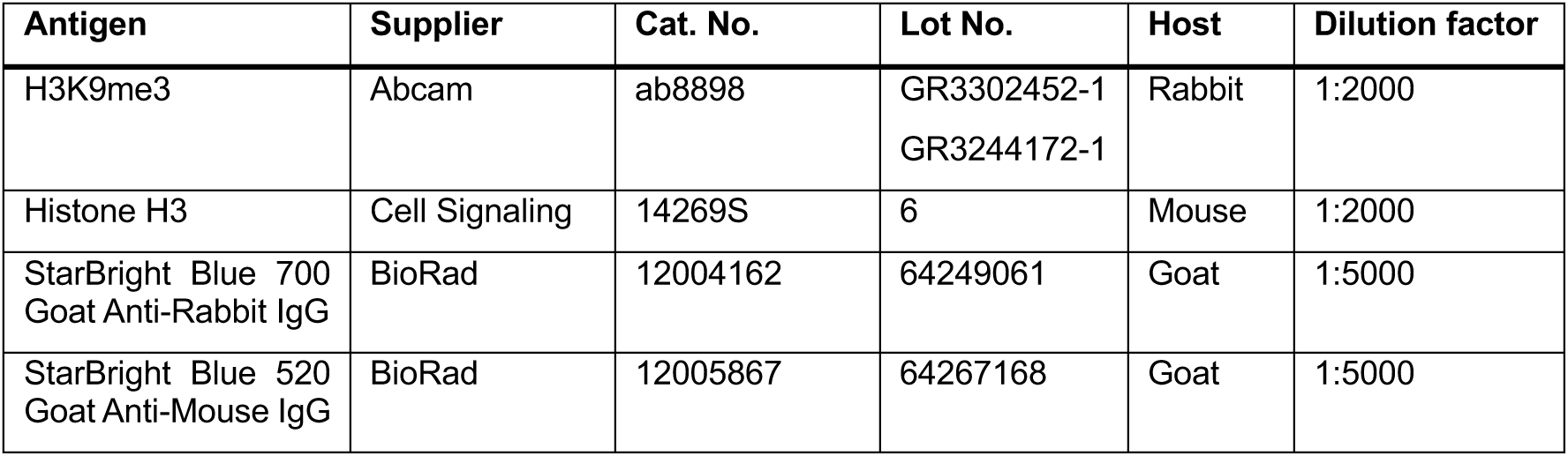
Antibody information for Western blot experiments.

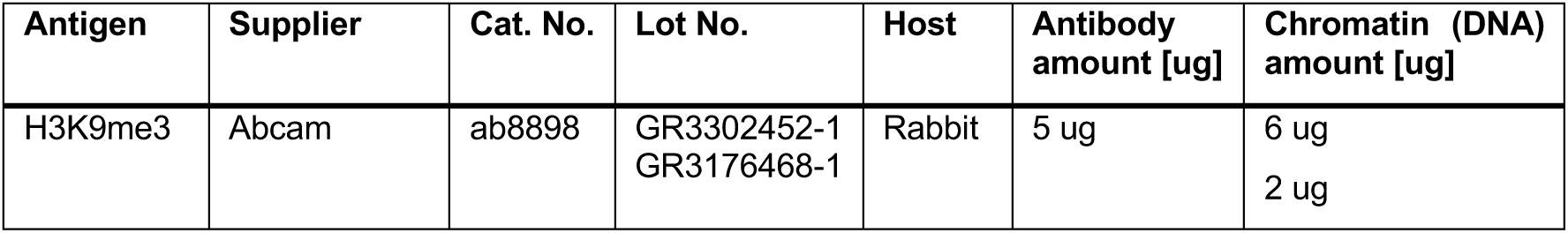
Antibody information for ChIP-seq experiments.

Antibody information for ChIP-seq experiments.
| Antigen | Supplier | Cat. No. | Lot No. | Host | Antibody amount [ug] | Chromatin (DNA) amount [ug] |
| --- | --- | --- | --- | --- | --- | --- |
| H3K9me3 | Abcam | ab8898 | GR3302452-1<br>GR3176468-1 | Rabbit | 5 ug | 6 ug<br>2 ug |

### Biological Resources

Human induced pluripotent stem cells (72.3, male, RRID: CVCL_A1BW)

### Statistical Analyses

Statistical tests, sample sizes, biological replicates, and error bars are described in the figure legend. Data were derived from individual differentiation of 72.3 hPSCs. Statistical analyses for Western blotting and imaging were performed using GraphPad Prism 9 and 10. A two-tailed unpaired t-test was used to compare differences between the two groups. For comparisons involving more than two groups, an Ordinary one-way ANOVA was applied. Detailed statistical analyses for RNA-seq and ChIP-seq, including differential gene expression, differential peak analysis, and Gene Ontology (GO) analysis, are provided in the method details.

### Human pluripotent stem cell (hPSC) culture

Human induced pluripotent stem cells (72.3, male, RRID: CVCL_A1BW) were obtained from the CCHMC Pluripotent Stem Cell Facility (PSCF) and cultured under feeder-free conditions in mTeSR1 medium (85850, StemCell Technologies) on plates coated with either hESC-qualified Matrigel (354277, Corning) or Cultrex Stem Cell Qualified Reduced Growth Factor Basement Membrane Extract (3434-010-02, Biotechne). hPSCs were cultured in a humidified incubator at 37℃ with 5% CO_2_.

Spontaneously differentiated cells were manually removed under a stereomicroscope before passaging and differentiation. For passaging, hPSC colonies were dissociated into small clusters using Gentle Cell Dissociation Reagent (07174, STEMCELL Technologies), seeded onto Matrigel-or Culturex-coated plates, and cultured in mTeSR1. The step-by-step culture protocol is detailed in STAR Protocols (38).

### Drosophila cell culture

Drosophila Schneider 2 (S2) cells were cultured in Schneider’s Medium (21720024, Thermo Fisher Scientific) supplemented with 10% fetal bovine serum (FBS) and 1x penicillin-streptomycin (15140122, Thermo Fisher Scientific) at room temperature under normoxic conditions. FBS was heat-inactivated at 56℃ for 30 min before use. For cryopreservation, S2 cells were suspended in freezing medium consisting of 45% S2-conditioned medium, 45% FBS, and 10% DMSO, and then stored at –80℃. The S2-conditioned medium was filtered using a 0.22 μm filter unit before use.

### hPSC differentiation into the three germ layers

**Basal Medium for paraxial mesoderm and definitive endoderm differentiation:** For paraxial mesodermal (PM) and definitive endodermal (DE) differentiation, CDM2 was used as the basement medium (39). CDM2 consists of a 1:1 mixture of GlutaMAX-containing IMDM (31980097, Thermo Fisher Scientific) and GlutaMAX-containing F12 (31765092, Thermo Fisher Scientific), supplemented with 1 mg/ml polyvinyl alcohol (363170, Sigma-Aldrich), 1%v/v chemically defined lipid concentrate (11905031, Thermo Fisher Scientific), 450 μM 1-thioglycolic (M6145, Sigma-Aldrich), 0.7 μg/ml Insulin (I0516-5ML, Sigma-Aldrich), and 15 μg/ml transferrin (354277, Sigma-Aldrich). CDM2 was sterile-filtrated using a 0.22 μm filter unit, aliquoted, and stored at –80℃. CDM2 aliquots were used within two months.

**Paraxial mesoderm (PM) differentiation:** PM differentiation was performed as previously described (39). hPSCs were plated on Matrigel-or Culturex SCQ-coated plates (dilution factor as per hPSC culture) one day before induction. For mesendoderm (ME) differentiation, hPSCs were cultured in CDM2 medium supplemented with 33 ng/ml ActivinA (800-0, Shenandoah; or GFH6, Cell Guidance Systems), 4 μM CHIR99021 (SML1046-25MG, Sigma-Aldrich), 20 ng/ml bFGF (PHG0261, Thermo Fisher Scientific), and 100 nM PIK90 (528117, EMD Millipore) for 24 hours. For subsequent PM differentiation, ME cells were briefly washed with DMEM/F12 and cultured in CDM2 medium supplemented with 1 μM A83-01 (SML0788-5MG, Sigma-Aldrich), 3 μM CHIR99021, 20 ng/ml bFGF, and 250 nM LDN193189 for 24 hours.

**Definitive endoderm (DE) differentiation:** DE differentiation was performed as previously described (20, 40). hPSCs were plated on Matrigel-or Culturex SCQ-coated plates (dilution factor as per hPSC culture) one day before induction. For ME induction, hPSCs were cultured in CDM2 medium supplemented with 100 ng/ml ActivinA, 2 μM CHIR99021, and 50 nM PI103 (SML0559-5MG, Sigma-Aldrich) for 24 hours. For subsequent DE differentiation, ME cells were briefly washed with DMEM/F12 and cultured in CDM2 medium supplemented with 100 ng/ml ActivinA and 250 nM LDN193189 for 24 hours.

**Neural ectoderm (NE) differentiation:** Neural ectoderm (NE) differentiation was performed as previously described (41). hPSCs were plated on Matrigel-(1:50 dilution) or Culturex SCQ-coated cell culture plates one day before differentiation. NE was induced by E6 medium (A1516401, Thermo Fisher Scientific) supplemented with 500 nM LDN193189 (SML0559-5MG, Sigma-Aldrich), 10 μM SB431542 (S4317-5MG, Sigma-Aldrich), and 5 μM XAV931 (X3004-5MG, Sigma-Aldrich). Cells were cultured in this medium for 48 hours with daily media changes.

### Inhibition of MEK/ERK signaling

Cells were treated with 1 μM of PD0325901 (MEK inhibitor; 72182, STEMCELL Technologies) or DMSO (vehicle control) for two days during differentiation from hPSC into NE, PM, and DE.

### Immunostaining

For all imaging experiments, cells were seeded on Thermanox coverslips (15 mm DIA, 72275-05, Electron Microscopy Sciences) inserted into a 24-well cell culture plate or CELL DESK LF (MS-9132, SUMITOMO BAKELITE CO, Ltd). Coating conditions were the same as described above. Cultured cells were briefly washed with DMEM/F12 and fixed with 4% paraformaldehyde (PFA) (15710, Electron Microscopy Sciences) in 1x PBS for 10 min at room temperature. Fixed cells were washed three times with washing buffer (1% BSA [A9647-50G, Sigma Aldrich] and 0.01% TritonX-100 in 1x PBS) for 5 min each and then permeabilized with 0.2% TritonX-100 in 1x PBS for 10 min. After permeabilization, cells were washed three times with washing buffer for 5 min each. Cells were then blocked with blocking buffer (5% BSA in 1x PBS) for 1 hour at room temperature and incubated with the diluted primary antibody (250 μL reaction volume) overnight at 4°C. The next day, cells were washed three times with washing buffer for 10 min each and then incubated with the diluted secondary antibody (250 μL reaction volume) for 2 hours at room temperature in the dark. After secondary antibody incubation, cells were washed three times with washing buffer for 10 min each. Nuclei were stained with 1 μg/mL DAPI (D9542-10MG, Sigma-Aldrich) in washing buffer for 5 min at room temperature, followed by two additional washes with washing buffer for 5 min each. For confocal imaging, stained samples were mounted with Fluoromount-G (0100-01, SouthernBiotech).

### Image acquisition

Conventional immunofluorescence images were acquired using the Nikon A1 and A1R systems (Nikon instrument) with the following objective lenses (Nikon instrument): Plan Apochromat Lambda 20x (Numerical Aperture = 0.75), CFI Apochromat Lambda S 40x water immersion (Numerical Aperture = 1.15), CFI Plan Apochromat Lambda D 40x (Numerical Aperture = 0.95), CFI Plan Apochromat IR 60x WI (Numerical Aperture = 1.27), and CFI Plan Apochromat VC 60XC WI (Numerical Aperture = 1.20). The pinhole size was automatically adjusted based on the longest fluorescence dye wavelength. Step size was optimized for each objective lens. All images were acquired at 512 x 512 or 1024 x 1024 pixels.

### Imaging data processing

1. Denoising: Acquired images were denoised using the “Denoise.ai” function in NIS Elements Advanced Research (version 5.20/5.30/5.42, Nikon Instrument). For manual denoising, the same parameters were applied to all images. Denoised images were then converted to *.ims format (specific to IMARIS) using IMARIS Converter (version 9.90 and 10.00, Bitplane).
2. Max projection image preparation: 2D images were generated using ImageJ Fiji (https://imagej.net/software/fiji/). Maximum intensity projection was applied to all 2D images.

### Western blot assay

For nuclear protein extraction, cell pellets were suspended in ice-cold Buffer A (10 mM HEPES-NaOH [pH 7.9], 10 mM KCl, 0.1 mM EDTA, 0.1M EGTA, 1 mM DTT, 1x Complete protease inhibitor EDTA-free in distilled water) and incubated on ice for 15 min. After incubation, 10% NP-40 was added to a final concentration of 0.625%, and the mixture was vortexed for 10 sec. The cell suspension was then centrifuged at 4,000 rpm for 5 min at 4℃ using an Eppendorf 5417R centrifuge. The supernatant was discarded, and the pellet was resuspended in 25 to 50 μL Buffer C (20 mM HEPES-NaOH [pH 7.9], 1 M NaCl, 1 mM EDTA, 1 mM EGTA, 1 mM DTT, 1x Complete protease inhibitor EDTA-free in distilled water). The suspension was incubated on ice for 15 min. Following nuclear lysis, the cell lysate was centrifuged at 10,000 rpm for 5 min at 4℃ using an Eppendorf 5417R centrifuge, and the supernatant was transferred to a new tube. Protein concentration was measured using the Bradford assay.

For Western blot analysis, 0.5 – 1.0 μg of nuclear protein in Laemmli Sample Buffer (J60015, Alfa Aesar) was denatured at 100℃ for 5 min, then separated on a 15% SDS-PAGE gel at 60 V for 30 min, followed by 100 V for 2 – 2.5 hours. Proteins on the gel were transferred to a 0.2 μm pore size nitrocellulose membrane (1620112, Bio-Rad) using a wet tank transfer system with the Mini Blot Module (Thermo Fisher Scientific) at 10 V for 1 hour. The transferred membrane was briefly washed with TTBS (0.05% Tween-20 in TBS) and blocked with blocking buffer (5% Blotting-Grade Blocker [1706404, Bio-Rad] in TBS) for 1 hour at room temperature. The blocked membrane was washed with TTBS at room temperature and incubated with the primary antibody, diluted in 5% BSA in TTBS, at 4℃ overnight. The next day, the membrane was washed three to five times with TTBS for 5 min each at room temperature, then incubated with the secondary antibody, diluted in blocking buffer, for 1 hour at room temperature. After incubation, the membrane was washed three to six times with TTBS for 5 min each at room temperature. Fluorescence imaging was performed using the ChemiDoc^TM^ MP imaging system (Bio-Rad).

### Spike-in control preparation

For ChIP-seq, crosslinked whole S2 cells were used as a spike-in control. S2 cells were crosslinked with 1% formaldehyde (F79-500, Fisher Scientific) in 1x PBS for 10 min at room temperature. The crosslinking reaction was quenched by adding 0.125 M glycine (15527013, Thermo Fisher Scientific) for 5 min at room temperature. The crosslinked cell suspension was centrifuged at 2,000 x g for 5 min at 4°C, and the supernatant was removed. The pellet was washed with ice-cold PBS and resuspended in 10% FBS in 1x PBS at a concentration of 2.0 or 4.0 x 10^7^ cells/mL. The suspension was aliquoted and centrifuged again at 2,000 x g for 5 min at 4°C. Finally, the cell pellet was frozen on dry ice and stored at –80°C until use.

### Chromatin Immunoprecipitation (ChIP)

The step-by-step ChIP protocol is detailed in STAR Protocols (42).

Cells were crosslinked with 1% formaldehyde in 1x PBS for 10 min at room temperature. The crosslinking reaction was quenched by adding 0.125 M glycine for 5 min at room temperature. Crosslinked cells were centrifuged at 600 x g for 5 min at 4°C, and the supernatant was removed. The cell pellet was washed with ice-cold 1x PBS, centrifuged again at 600 x g for 5 min at 4°C, and the supernatant was aspirated. This wash step was repeated one more time. The final cell pellet was frozen on dry ice and stored at –80°C until use.

Crosslinked human cells were resuspended in Lysis Buffer 1 (10 mM Tris-HCl [pH 8.0], 10 mM NaCl, 0.5% NP-40, and 1x Complete Protease Inhibitor EDTA-free) and incubated on ice for 10 min. For Drosophila S2 spike-in controls, crosslinked S2 cells were resuspended in 100 μL Lysis Buffer 1 at a final concentration of 2.0 x 10^5^ or 4.0 x 10^5^ cells/μL, and 25% of the S2 cells were mixed with human cells. The mixed cell suspension was centrifuged at 665 x g for 5 min at 4℃, and the supernatant was removed. The cell pellet was then resuspended in Lysis Buffer 2 (50 mM Tris-HCl [pH 8.0], 10 mM EDTA, 0.32% SDS, 1x Complete Protease Inhibitor EDTA-free) and incubated on ice for 10 min. The cell lysate was then diluted in IP Dilution Buffer (20 mM Tris-HCl [pH 8.0], 150 mM NaCl, 2 mM EDTA, 1% TritonX-100, 1x Complete Protease Inhibitor EDTA-free) and transferred to a milliTUBE ATA Fiber (Covaris). Chromatin was sonicated using a Covaris S220 (Covaris) for 5.5 min. Insoluble debris was removed by centrifugation at 12,000 x g for 5 min at 4℃. The supernatant containing sonicated chromatin was transferred to a new tube and stored at –80℃ until use.

Dynabeads Protein-G (10004D, Thermo Fisher Scientific) (25 – 80 μL) were washed twice with PBS-T (0.02% Tween-20 in 1x PBS). The desired amounts of antibodies were conjugated to washed Dynabeads by incubating for 2 to 6 hours at 4°C with rotation. Meanwhile, frozen sonicated chromatin was thawed on ice. A desired amount of chromatin was diluted with IP dilution buffer at a 4:1 ratio. Antibody-conjugated Dynabeads were washed twice with IP Dilution Buffer (without protease inhibitor), resuspended in the diluted chromatin, and incubated overnight at 4°C with rotation. The next day, ChIPed beads were washed four times using the following washing buffers: (1) FA Lysis Buffer (50 mM HEPES-KOH [pH 7.5], 150 mM NaCl, 2 mM EDTA, 1% TritonX-100, 0.1% Sodium deoxycholate, 1x Complete protease inhibitor EDTA-free); (2) NaCl Buffer (50 mM HEPES-KOH [pH 7.5], 500 mM NaCl, 2 mM EDTA, 1% TritonX-100, 0.1% Sodium deoxycholate); (3) LiCl Buffer (100 mM Tris-HCl [pH 8.0], 500 mM LiCl, 1% NP-40, 1% Sodium deoxycholate); (4) 10 mM Tris-HCl [pH 8.0]. Following washing steps, chromatin was eluted with TES buffer (50 mM Tris-HCl [pH 8.0], 10 mM EDTA, 1% SDS) for 15 min at 65°C while shaking at 1,000 rpm in ThermoMixer F1.5 (Eppendorf). The ChIP and Input chromatin were reverse-crosslinked by adding NaCl to a final concentration of 200 mM and incubating for 8 hours at 65°C. Samples were then treated with 50 ng/uL RNaseA (EN0531, ThermoFisher Scientific) for 30 min at 37°C, followed by treatment with 0.2 mg/mL Protease K (03115828001, Roche) for 2 hours at 37°C. DNA was purified using phenol-chloroform extraction, followed by ethanol precipitation. DNA concentration was measured using a Quantus fluorometer (Promega).

### Preparation and sequencing of ChIP-seq libraries

Multiplexed libraries were prepared from ChIP and Input samples (two replicates) using the NEBNext Ultra II DNA Library Prep Kit for Illumina (E7645, NEB), following the manufacturer’s protocol with the following modifications. After adaptor ligation, cleanup was performed using AMPure XP Beads (A63880, Beckman Coulter), MagBind Magnetic Beads (M1378-01, Omega), or homemade Sera-mag beads (43) without size selection. After RCR enrichment of the adaptor-ligated library, two rounds of size selection were performed using the AMPure XP Beads or MagBind Magnetic Beads. Paired-end 38 bp or 50 bp sequencing was performed on an Illumina NextSeq500 or NovaSeqX Plus.

### Preparation and sequencing of RNA-seq libraries

1.0×10^6^ cultured cells were dissociated into single cells using Accutase and were mixed with 100,000 *Drosophila* Schneider 2 (S2) cells for a spike-in control. Cells were lysed with Aurum Total RNA Lysis Solution (7326802, BioRad). Total RNA was extracted using Aurum™ Total RNA Mini Kit (7326820, BioRad) and sent to Novogene for library preparation and paired-end 150 bp sequencing on an Illumina NovaSeq 6000 or NovaSeq X Plus.

### RNA-seq data analysis

RNA-seq reads were aligned to the UCSC human genome hg38 using the STAR aligner (44) to match the analysis environment with our previously published data (31). Only uniquely aligned reads were retained for the downstream analysis. Read counts were measured based on the UCSC RefSeq gene annotations of hg38. Differential gene expression analysis using DESeq2 (45). Differentially expressed genes were identified by FDR < 0.01 and fold-change > 2. Gene ontology analysis was performed using Enrichr (46). Heatmaps were generated by R Studio with the gplots and genefilter packages.

### ChIP-seq data analysis

First, we created a custom reference by combining the UCSC human genome hg38 and the *Drosophila* genome dm6. ChIP-seq reads were aligned to the custom reference combining hg38 and dm6 using the STAR aligner. Only uniquely aligned reads were retained for downstream analysis. Differential H3K9me3 domain analysis was performed in multiple steps, combining Homer (47) peak calling and differential analysis using DESeq2 to leverage the statistical power of biological replicates. First, preliminary variable-width peak calling is performed for each ChIP replicate without an input control. Next, peaks from replicates were pooled and merged from biological replicates for each condition to prepare peak candidates. 2 kb-sized tiles were prepared with the peak candidates sliding by 1kb for differential analysis comparing ChIP and Input. Differential analysis was performed for tiles using DESeq2 without considering spike-in control and differential tiles (FC > 2.0 & p-value < 0.01 for H3K9me3). ChIP-specific tiles were stitched together if their distance was < 4kb as a final peak set for each condition. Differential domain analysis between conditions was performed similarly by DESeq2, but incorporating spike-in reads this time. Spike-in read counts were measured for each replicate, which was then used to calculate the sizeFactor for a DESeqDataSet object. Each set of condition-specific differential tiles was identified between conditions (FC > 1.5 & p-value < 0.01 for H3K9me3) and stitched together to make the final differential domains if their distance was < 4kb. Differential domains were identified in association with nearby genes using GREAT (48). Spearman’s correlation coefficient-based clustering was performed with the plotCorrelation in deepTools (49). All browser views were generated with the UCSC genome browser. Lastly, spike-in normalized ChIP tracks were generated for visualization in a genome browser track. Instead of a conventional read-per-million scale, we used spike-in read count and normalized the human reads in read-per-0.1-million spike-in count.

## Results

### H3K9me3 heterochromatin undergoes global reorganization during mesoderm and endoderm differentiation in association with EMT

To examine how H3K9me3 domains are rearranged along with gene expression changes during human germ layer differentiation, we employed established hPSC differentiation protocols for neural ectoderm (NE) (41), paraxial mesoderm (PM) (39), and definitive endoderm (DE) (40), which mimic the stepwise activation and inhibition of signaling pathways during development (**Figure 1A**). Successful differentiation was validated through immunofluorescence for lineage-specific markers (POU5F1/OCT4 and SOX2 for hPSC; SOX1 and SOX2 for NE; TBX6 and CDX2 for PM; SOX17 and FOXA2 for DE) (**Figure S1A**). RNA-seq further confirmed lineage-specific marker expression and gene ontology (GO) terms enrichment in NE (e.g., *SOX1*, *HES5*, and *FOXG1;* “Nervous System Development”), PM (e.g., *TBX6*, *CDX2*, *HES7*, and *MSGN1;* “Skeletal System Development”), and DE (e.g., *SOX17*, *FOXA2*, *GATA6*, and *CER1;* “Endoderm formation”) (**Figure S1B, S1C**). We found that differentiation of hPSCs into PM and DE induced a greater number of differentially expressed genes than differentiation into NE (**Figure S1C)**. Notably, principal component analysis (PCA) of RNA-seq data revealed three distinct clusters: while hPSC and NE clustered together, PM and DE formed clearly separated clusters (**Figure 1B**). This clustering pattern reflects differences in whether differentiated cells are undergoing EMT or not. Indeed, we confirmed that PM and DE cells homogenously expressed mesenchymal cadherin, CDH2, whereas hPSC and NE cells predominantly expressed epithelial cadherin, CDH1 (**Figure 1C**). To further validate EMT-related cytoskeleton features, we stained filamentous actin (F-actin) with Phalloidin. In hPSCs and NE cells, F-actin was predominantly localized at cell-cell adhesions and formed cobblestone-like structures, hallmarks of epithelia. In contrast, in PM and DE cells, F-actin was assembled into stress fibers and formed mesenchymal cell-like stretched structures (**Figure S1D**). These results confirm that PM and DE acquire mesenchyme-like characteristics, while NE retains epithelial traits.

**Figure 1.**
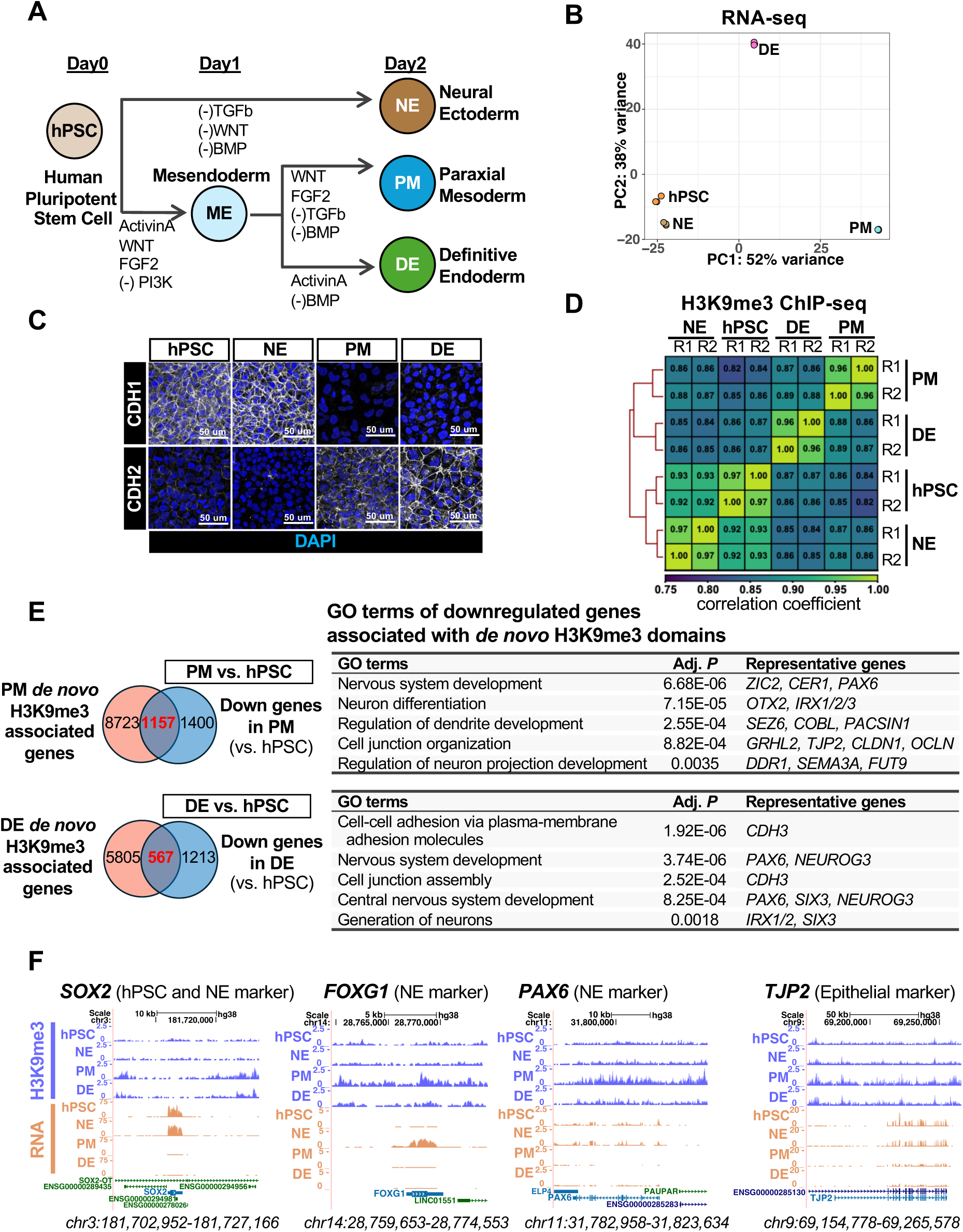
H3K9me3 heterochromatin undergoes global reorganization during mesoderm and endoderm differentiation in association with EMT. (**A**) Schematic overview of the differentiation protocols utilized to generate neural ectoderm (NE), paraxial mesoderm (PM, and definitive endoderm (DE) from human pluripotent stem cells (hPSCs). **(B)** Principal component analysis (PCA) of RNA-seq data from hPSC, NE, PM, and DE, based on the top 3,000 expressed genes (n = 3 replicates from three independent differentiations). **(C)** Immunostaining of CDH1 (epithelial marker) and CDH2 (mesenchymal marker), along with nuclear staining (DAPI), in hPSC and germ layers. Scale bars, 50 μm. **(D)** Spearman correlation analysis of H3K9me3 ChIP-seq data across individual replicates of hPSC and germ layers. (n = 2 replicates from two independent differentiations). **(E)** Intersection between genes associated with *de novo* H3K9me3 domains and genes downregulated during hPSC differentiation into PM and DE. GO terms associated with these downregulated genes are shown, along with representative genes. Adjusted *P*-values were calculated using Enrichr. **(F)** Genome browser tracks displaying H3K9me3 ChIP-seq and RNA-seq data for hPSC and germ layers at representative loci of hPSC, NE, and epithelial marker genes.

To investigate the genome-wide localization of H3K9me3 domains during germ layer differentiation, we performed chromatin immunoprecipitation sequencing (ChIP-seq). To appropriately normalize the global enrichment of H3K9me3 marks, we included *Drosophila* S2 cells as a spike-in control (50). Differential H3K9me3 domain analysis between hPSCs and the germ layers indicated a greater number of newly established, *de novo* (gained) H3K9me3 domains in PM (n =16,903 domains) and DE (n = 8,643 domains), compared to NE (n = 2,922 domains) (**Figure S2A**). These results are consistent with a previous report that the number of genes marked by H3K9me3 is increased from mouse epiblast to mesoderm and endoderm differentiation (34). Western blot assays also confirmed that the bulk abundance of H3K9me3 relative to H3 was increased during hPSC differentiation into PM and DE, but not NE (**Figure S2B**). Notably, genome-wide Spearman’s rank correlation analysis of the H3K9me3 ChIP-seq signals revealed that hPSC/NE, PM, and DE form distinct clusters (**Figure 1D**), similar to the clustering pattern observed in RNA-seq analysis (**Figure 1B**). These results suggest that the lineage-specific H3K9me3 landscape is reorganized during PM and DE differentiation, along with EMT occurring, and concurrent with dramatic gene expression changes.

To characterize target genes of the newly established, *de novo* H3K9me3 domains in PM and DE, we first associated these domains with potential target genes using the Genomic Regions Enrichment of Annotations Tool (GREAT) (48). We then intersected the annotated genes with those downregulated in PM and DE compared to hPSC and the other lineages (**Figure 1E, S2C**). Subsequently, we performed gene ontology (GO) analysis to functionally characterize potential targets of *de novo* H3K9me3-mediated repression. We found that those genes in PM and DE were characterized by GO terms related to neural development, including “Nervous system development” and “Cell junction assembly” (**Figure 1E, S2C**). For instance, key NE markers (*SOX2*, *FOXG1*, *PAX6*) and an epithelial marker (*TJP2*) were downregulated and associated with *de novo* H3K9me3 domains in PM and DE (**Figure 1F**). These results suggest that the H3K9me3 reorganization contributes to repressing alternative, neural lineage genes in PM and DE differentiation.

### Ectopic H3K9me3 domain formation occurs in the absence of MAPK/ERK signaling during mesoderm and endoderm differentiation

While FGF-mediated MAPK/ERK signaling pathway has been shown to regulate EMT and the specification of mesoderm and endoderm during gastrulation (12–19), its underlying mechanisms remain less well understood amongst the key gastrulation signals. To investigate whether MAPK/ERK pathway may have a role in H3K9me3 reorganization during germ layer differentiation, we inhibited MEK with the PD0325901 inhibitor (MEKi) (18, 51) during PM and DE differentiation, as well as NE differentiation as a control (**Figure 2A**). We first validated the effects of MEKi on lineage marker expression and EMT states using immunostaining. As expected, MEKi treatment in PM and DE resulted in the loss of lineage marker expression (TBX6 in PM and SOX17 in DE) and the upregulation of NE markers (SOX1 and SOX2) (**Figure 2B**). In contrast, MEKi treatment in NE did not affect NE marker expression (SOX1 and SOX2) (**Figure S3A**). We also confirmed that MEKi treatment compromised EMT in PM and DE, leading to an upregulation of the epithelial marker CDH1 (**Figure 2C**) and the typical epithelial-like cell morphology with F-actin structures (**Figure S3B**). As expected, MEKi treatment did not alter the epithelial features of NE (**Figure S3B-C**). These results confirm that the MAPK/ERK signaling pathway regulates EMT and promotes mesoderm and endoderm specification.

**Figure 2.**
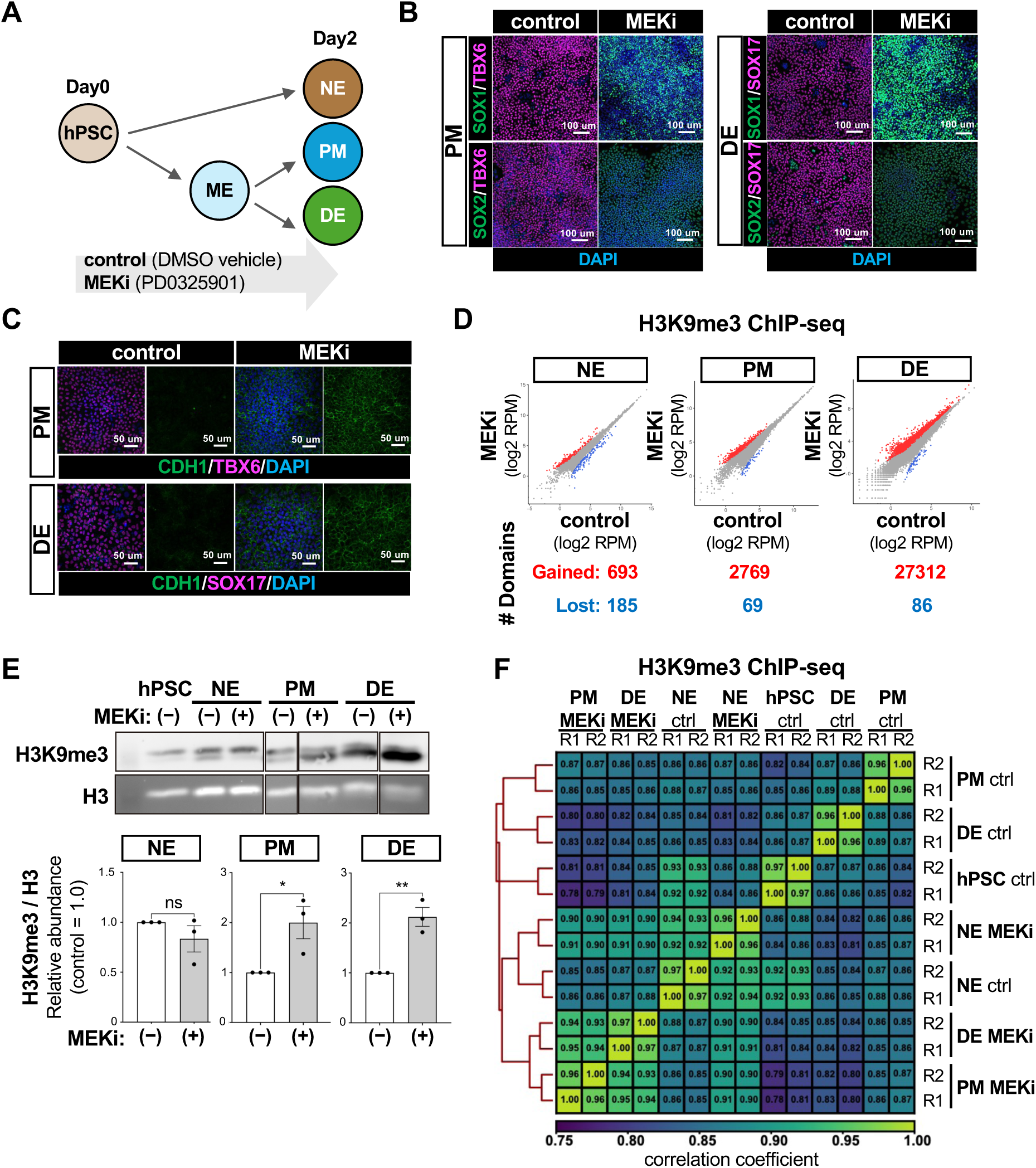
Ectopic H3K9me3 domain formation occurs in the absence of MAPK/ERK signaling during mesoderm and endoderm differentiation. **(A)** Schematic overview of MEK inhibition using PD0325901 (MEKi) during hPSC differentiation into germ layers. **(B)** Immunostaining of the NE markers (SOX1 and SOX2), PM marker (TBX6), and DE marker (SOX17), along with nuclear staining (DAPI), in control and MEKi-treated PM and DE. Scale bars,100 μm. **(C)** Immunostaining of epithelial marker (CDH1) and lineage markers (TBX6 for PM, SOX17 for DE), along with nuclear staining (DAPI), in control and MEKi-treated PM and DE. Scale bars, 50 μm. **(D)** Differential H3K9me3 domain analysis comparing control and MEKi-treated cells (n = 2 replicates from two independent differentiations; *P* < 0.01 by DESeq2). **(E)** Western blot quantification of relative H3K9me3 signal intensity, normalized to total H3 (n = 3 replicates from three independent differentiations, means ± SEM; \**P* < 0.05, \*\**P* < 0.02, ns = not significant, Student’s *t*-test). **(F)** Genome-wide Spearman correlation analysis of H3K9me3 ChIP-seq data across individual replicates of control and MEKi-treated cells.

To examine if MAPK/ERK signaling impacts H3K9me3 reorganization during PM and DE differentiation, we performed H3K9me3 ChIP-seq on MEKi-treated cells. Strikingly, MEKi treatment led to a marked increase in the number of gained H3K9me3 domains, particularly in PM (n=2,769) and DE (n=27,312), compared to NE (n = 693) (**Figure 2D**). To validate the global increase of H3K9me3 domains in PM and DE, we measured the abundance of H3K9me3 modifications relative to bulk H3 by Western blotting analysis. Consistent with the ChIP-seq results, PM and DE, but not NE, showed a significant increase in H3K9me3 modifications (**Figure 2E**). The expression of both H3K9 methyltransferases and demethylases was moderately upregulated in MEKi-treated cells (**Figure S3D**). Therefore, the global increase in H3K9me3 modifications could not be solely explained by changes in the H3K9 modifier expression. Spearman’s rank correlation analysis of the H3K9me3 ChIP-seq data revealed that MEKi treatment altered the H3K9me3 distribution pattern in PM and DE, making it more similar to NE (**Figure 2F**). These findings suggest that MAPK/ERK signaling prevents ectopic H3K9me3 domain formation in PM and DE and is required for the global reorganization of the H3K9me3 landscape during germ layer differentiation.

### MAPK/ERK signaling regulates EMT and lineage-specific gene expression in mesoderm and endoderm differentiation

Having shown that MAPK/ERK signaling prevents ectopic H3K9me3 domain formation specifically during human mesoderm and endoderm differentiation, we aimed to identify the gene loci regulated via this pathway. As a first step, we performed RNA-seq on PM and DE cells treated with the MEKi during differentiation. Principal component analysis of the RNA-seq data revealed that MEKi treatment of PM and DE altered gene expression patterns more closely resembling those of NE (**Figure 3A**), consistent with the clustering pattern observed in the H3K9me3 ChIP-seq analysis (**Figure 2F**). Accordingly, upregulated genes in MEKi-treated PM and DE were characterized by GO terms related to neural lineages, such as “Axonogenesis” and “Nervous System Development” (**Figure 3B**). In contrast, downregulated genes in MEKi-treated PM and DE were associated with mesoderm-specific GO terms (e.g., “Aortic Valve Development”) and endoderm-specific GO terms (e.g., “Endoderm Formation”), respectively (**Figure 3B**). Key mesoderm regulators (e.g., *DLL3, TBX3, TBXT,* and *HES7*) and endoderm regulators (e.g., *SOX17, EOMES, GATA6,* and *HHEX*) were significantly downregulated following MEKi treatment (**Figure 3C**). Furthermore, EMT-related GO terms, including “Regulation of cell migration” and “Regulation of epithelial to mesenchymal transition,” were enriched among the downregulated genes in MEKi-treated PM and DE (**Figure 3B**). Notably, these included critical mesenchymal markers such as *CDH2* and EMT inducers like *TWIST2, ZEB1*, and *SNAI1* (**Figure 3C**). Taken together, these genome-wide analyses demonstrate that MAPK/ERK signaling is essential for regulating the expression of key EMT-related and lineage-specific genes in mesoderm and endoderm differentiation.

**Figure 3.**
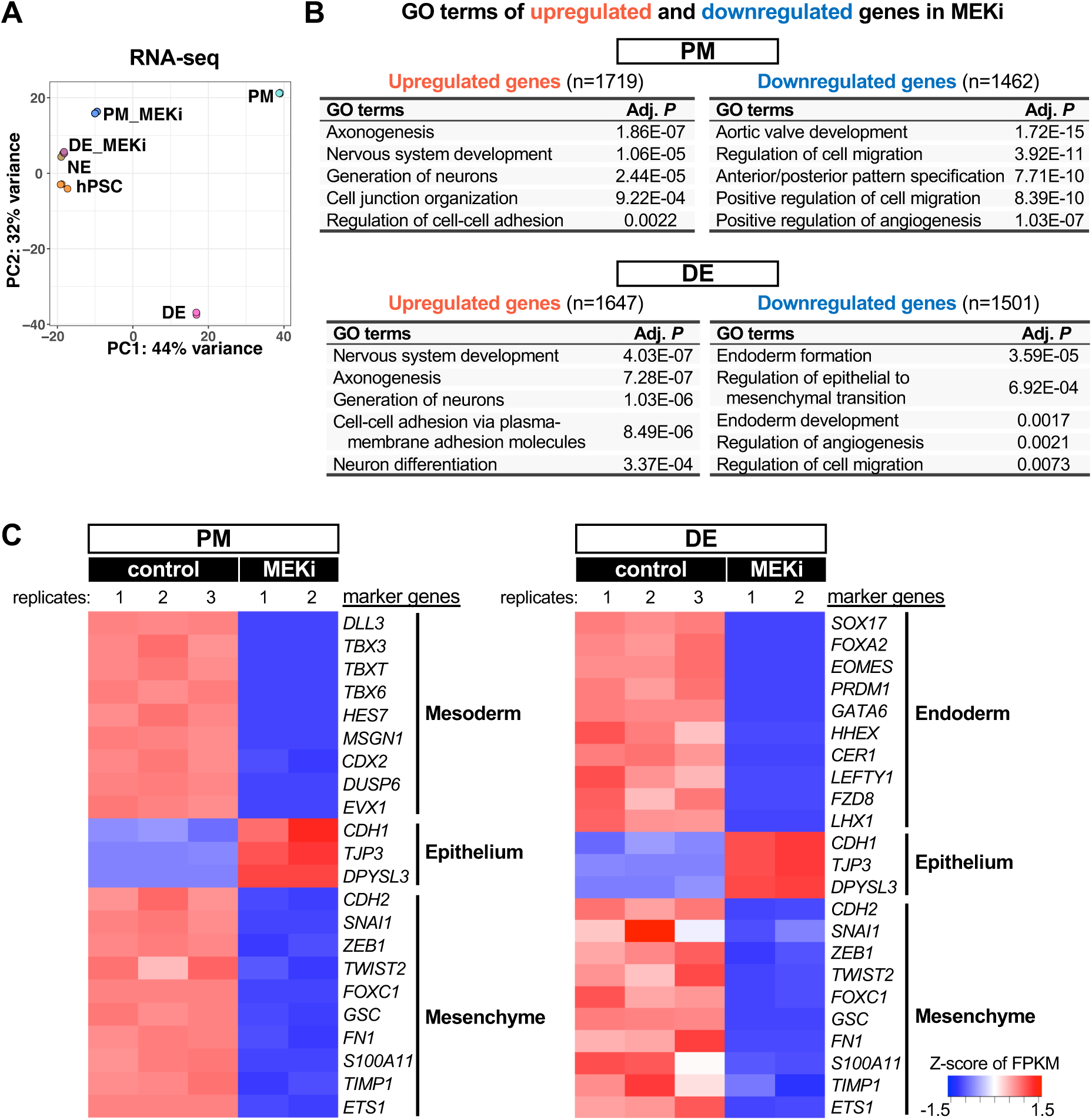
MAPK/ERK signaling regulates EMT and lineage-specific gene expression in mesoderm and endoderm differentiation. **(A)** Principal component analysis (PCA) of RNA-seq data from hPSC, NE, PM, DE, and MEKi-treated PM and DE, based on the top 3,000 expressed genes (n = 3 replicates from three independent differentiations for control cells, n = 2 replicates from two independent differentiations for MEKi-treated cells). **(B)** GO terms associated with upregulated (red) and downregulated (blue) genes in MEKi-treated PM and DE. **(C)** RNA-seq heatmaps of representative marker gene expression across control and MEKi-treated PM and DE in Z-score.

### MAPK/ERK signaling blocks the establishment of H3K9me3 heterochromatin domains at EMT– and lineage-specific gene loci necessary for mesoderm and endoderm differentiation

Having identified the genes whose expression was regulated by MAPK/ERK signaling during endoderm and mesoderm differentiation, we next examined how MAPK/ERK-mediated H3K9me3 reorganization impacts this lineage-specific gene expression. To identify genes potentially affected by ectopic H3K9me3 domain formation in MEKi-treated PM and DE, we first associated the gained H3K9me3 domains with potential target genes using the GREAT. We then intersected the annotated genes with those downregulated following MEKi treatment (**Figure 4A-B**). In PM, such genes included key mesodermal and EMT regulators, such as *TBXT*, *TBX3, DLL3,* and *TWIST2*, and were characterized by mesoderm-related GO terms, including “Skeletal system development” (**Figure 4A, 4C, S4A**). Similarly, in DE, genes with ectopically formed H3K9me3 domains and downregulated upon MEKi treatment included crucial endodermal and EMT regulators, such as *EOMES, GATA6, SOX17, HHEX,* and *TGFB2*, and were characterized by GO terms related to endoderm and EMT, including “Endoderm formation” and “Regulation of cell migration” (**Figure 4B, 4C, S4A**). Notably, all genes represented in the GO terms shown in **Figure 4A-B** play critical roles in mesoderm and endoderm differentiation, respectively. These results suggest that MAPK/ERK signaling prevents ectopic H3K9me3 domain formation at EMT and lineage-specific genes during mesoderm and endoderm differentiation.

**Figure 4.**
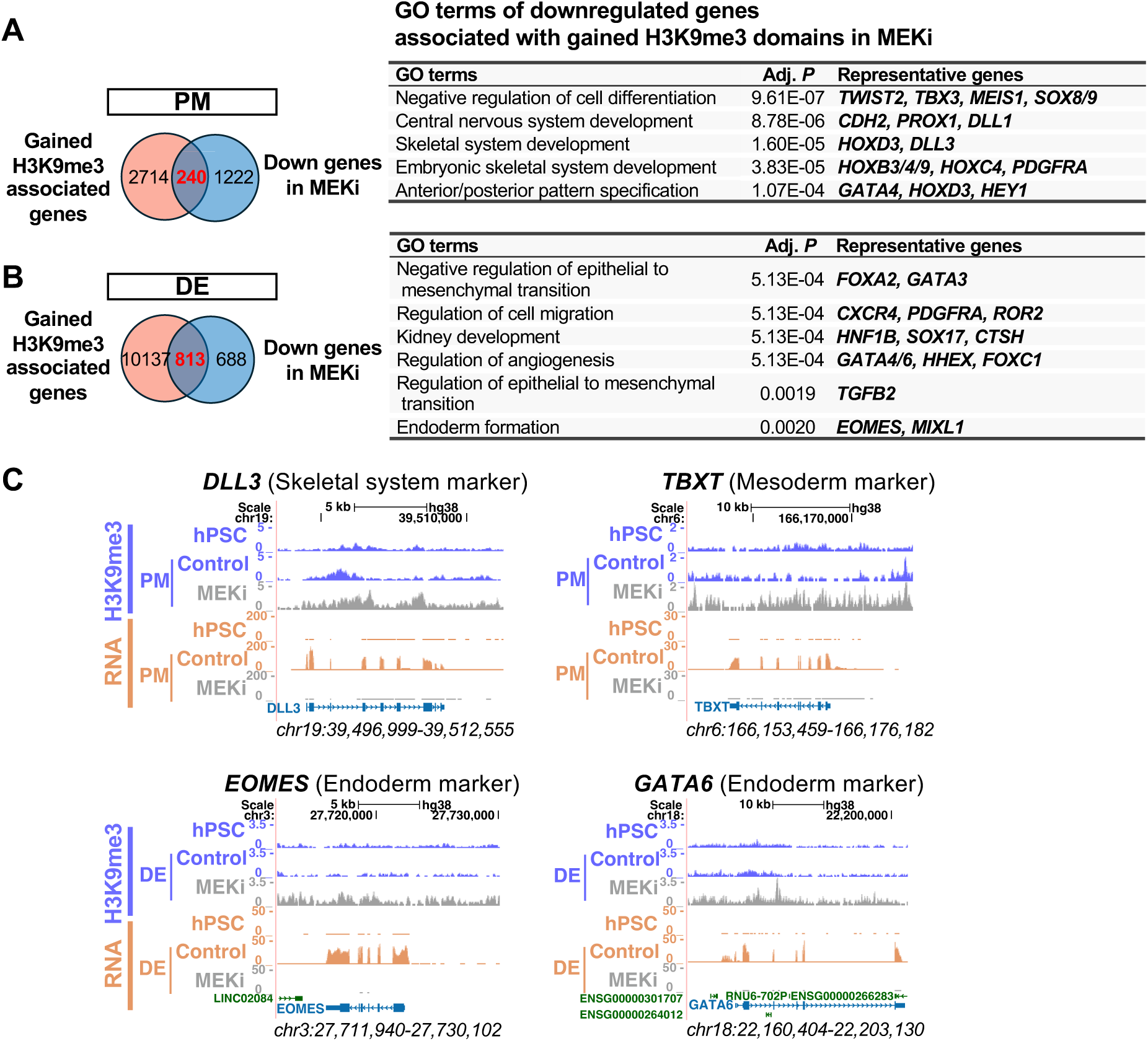
MEK/ERK signaling blocks the establishment of H3K9me3 domains at EMT– and lineage-specific gene loci necessary for mesoderm and endoderm differentiation. (**A**) Intersection between genes associated with gained H3K9me3 domains and genes downregulated in MEKi-treated PM. GO terms associated with these downregulated genes are shown, along with representative genes. Adjusted *P*-values were calculated using Enrichr. **(B)** Intersection between genes associated with gained H3K9me3 domains and genes downregulated in MEKi-treated DE. GO terms associated with these downregulated genes are shown, along with representative genes. Adjusted *P*-values were calculated using Enrichr. **(C)** Genome browser tracks displaying H3K9me3 ChIP-seq and RNA-seq data in PM and DE at representative loci of PM– and DE-related genes.

We further explored whether genes that were silent in both control and MEKi-treated cells were impacted by the ectopic formation of H3K9me3 domains upon MEKi treatment, as these genes may remain silenced but normally devoid of H3K9me3 formation for activation during later stages of differentiation. We intersected genes that were silent in both control and MEKi with ectopically formed H3K9me3 domains upon MEKi treatment (**Figure S4B-C)**. Representative genes fulfilling these criteria, shown in **Figure S4B-C,** play an important role in later mesodermal and endodermal tissue differentiation or functions. For instance, FOXA1 and PROX1 regulate liver and pancreas development, and PAX6 and NEUROG3 are essential for pancreatic development (52, 53). HGF (hepatocyte growth factor), WNT6/7B/11, and SHH contribute to endodermal tissue patterning and organogenesis (52, 53). Additionally, genes associated with the GO term “Chemical synaptic transmission” are involved in endoderm-derived organ functions, such as those of the pancreas and gastrointestinal tract, including SST (Somatostatin) (54, 55).

Collectively, MAPK/ERK signaling blocks the establishment of H3K9me3 domains at genes critical for EMT, as well as genes involved at early and later developmental stages in mesodermal and endodermal tissues. This highlights the critical role of MAPK/ERK signaling in maintaining the developmental plasticity necessary for proper differentiation and functions.

## DISCUSSION

During gastrulation, dynamic interplay among key cell signaling pathways, BMP, WNT, NODAL, and FGF, dictates cell fate decisions. While extensive studies have elucidated their critical roles in transcriptional and morphological regulation, how these signals orchestrate the epigenome to control cell differentiation remains largely unknown. In this study, we address this critical gap and reveal that the MAPK/ERK, a key downstream pathway of FGF signaling during gastrulation, reshapes the lineage-specific H3K9me3 landscape to enable proper mesoderm and endoderm specification.

Among the major signaling pathways governing gastrulation, the functional mechanisms of FGF signaling remain the least understood. Our findings mechanistically explain previous observations. For instance, it has been shown that NODAL signaling is insufficient to induce mesendoderm and endoderm formation without FGF-ERK signaling (17). Our results suggest that FGF/ERK signaling enables NODAL function through blocking ectopic H3K9me3 formation at its target genes. Similarly, in *Fgfr1* knockout mouse embryos, although *Wnt3a* expression is maintained during gastrulation, direct Wnt target genes containing TCF binding sites, such as *Tbxt,* fail to be induced (13). In addition to the previously proposed mechanism in which *Cdh1* derepression in *Fgfr1* mutants sequesters free Ctnnb1(13), our study suggests that *Fgfr1* mutants may ectopically establish H3K9me3 domains at Wnt target gene loci, thereby attenuating Wnt signaling transduction. Consistently, MEK inhibition in PM resulted in ectopic H3K9me3 accumulation at the *TBXT* gene locus and prevented *TBXT* induction (**Figure 4C**). Collectively, our study reveals a new role for MAPK/ERK signaling in facilitating the activity of other signaling pathways, such as NODAL and WNT, by blocking H3K9me3-mediated gene repression at key lineage regulators.

An unaddressed question that arises is how MAPK/ERK signaling prevents ectopic H3K9me3 heterochromatin formation. In *C. elegans,* global H3K9me3 domains increase as cells progress toward gastrulation (56, 57). Consistently, we observed a global increase in H3K9me3 domains during hPSC differentiation into PM and DE, but not NE differentiation (**Figure S2A-B**). Our findings suggest that during this global H3K9me3 expansion, MAPK/ERK signaling actively counteracts H3K9me3 formation at EMT-, mesoderm-, and endoderm-specific gene loci. The MAPK/ERK signaling cascade functions through sequential phosphorylation and activation of protein kinases and influences multiple cellular processes, including transcriptional regulation. Given that over 160 MAPK/ERK substrates have been reported (58–60), identifying the downstream MAPK effectors that block H3K9me3 formation remains challenging. However, transcription factors and chromatin-related factors are likely candidates, as active chromatin-associated mechanisms have been proposed to protect genes from heterochromatin invasion (61, 62). For instance, in yeast, Set1/COMPASS-mediated H3K4 methylation repels heterochromatin invasion by inhibiting H3K9 methyltransferase Suv39/Clr4 activity and locally disrupting nucleosome stability (63). Identifying the MAPK/ERK downstream effectors that prevent H3K9me3 invasion will further advance our understanding of how this pathway regulates gene expression, not only during gastrulation but also across broader developmental and pathological contexts.

## Data Availability

Sequencing data are available through GEO accession numbers GSE215436 (RNA-seq in hPSC and DE), GSE289666 (ChIP-seq), and GSE289669 (RNA-seq).

## Acknowledgments

We thank D. Haslam, B. Gebelein, A. Barski, J. Tchieu, and T. Takebe for sharing equipment and materials; J. Tchieu and K. Loh for technical advice for cell differentiation protocols; N. Bushati for helpful comments and editing on the paper; and the Digestive Disease Research Core Center in CCHMC (Pluripotent Stem Cell Facility, DNA sequencing Core, Bio-Imaging and Analysis Facility).

## Author Contributions

Conceptualization: S.M., M.I.

Investigation: S.M., M.I., M.G., S.S., G.M.

Formal analysis: S.M., H-W.L.

Software: H-W.L., S.M., R.M., M.T., C.A.

Visualization: S.M., M.I.

Writing – Original Draft: M.I., S.M.

Writing – Review & Editing: All authors

Supervision and Funding acquisition: M.I.

---

**Figure S1.**
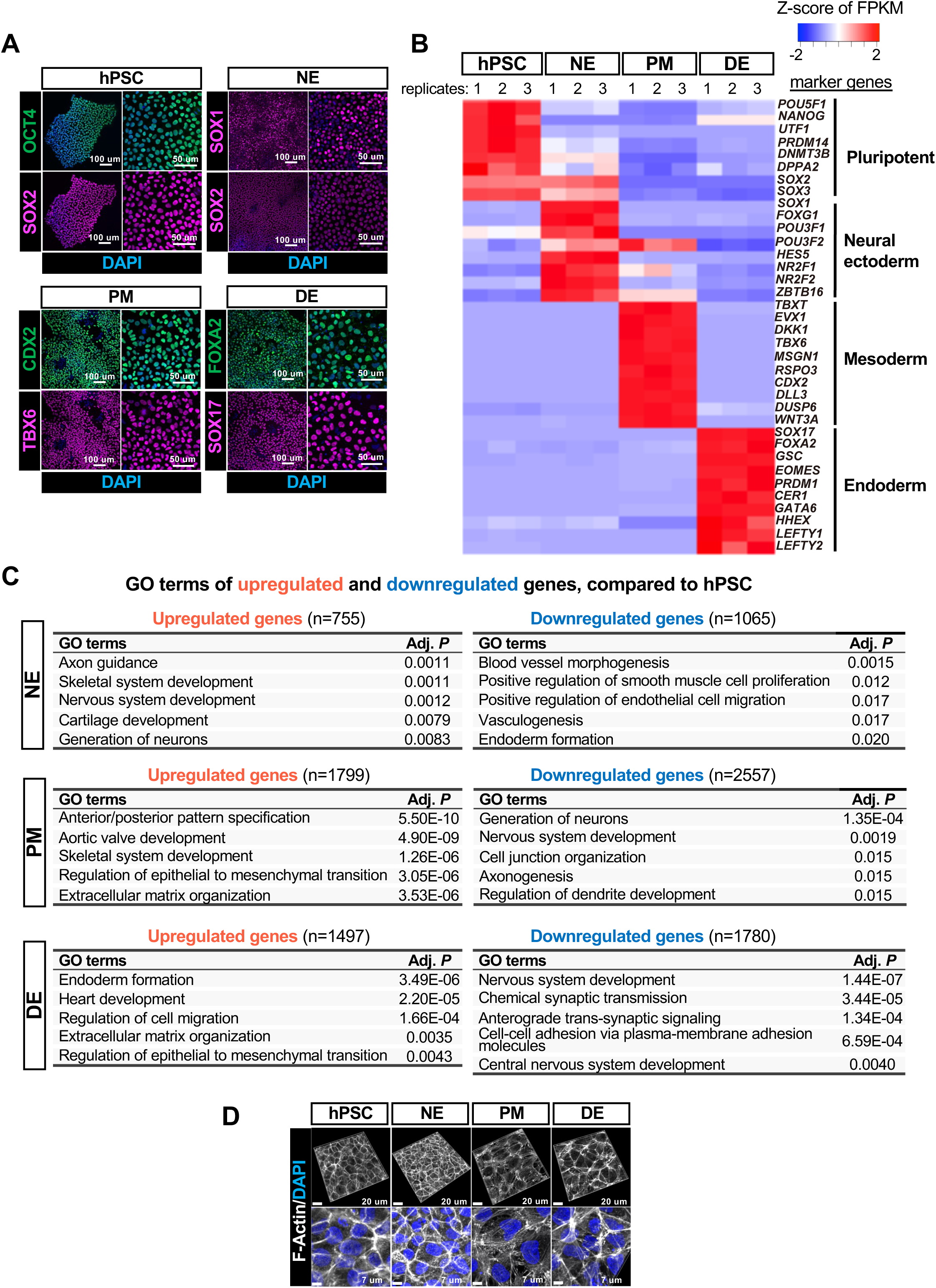
Characterization of hPSCs and germ layers, Related to Figure S1. **(A)** Immunostaining showing expression of lineage-specific transcription factors. Lower (left) and higher (right) magnification are shown. Scale bars, 100 μm (left) and 50 μm (right). **(B)** RNA-seq heatmap of representative marker gene expression across hPSCs and germ layers in Z-score (n = 3 replicates from three independent differentiations). **(C)** Gene ontology (GO) terms associated with upregulated (red) and downregulated (blue) genes during hPSC differentiation into germ layers. Adjusted *P*-values were calculated using Enrichr. **(D)** 2D and 3D visualization of F-actin patterns in hPSCs and germ layers. Scale bars, 20 μm (top) and 7 μm (bottom).

**Figure S2.**
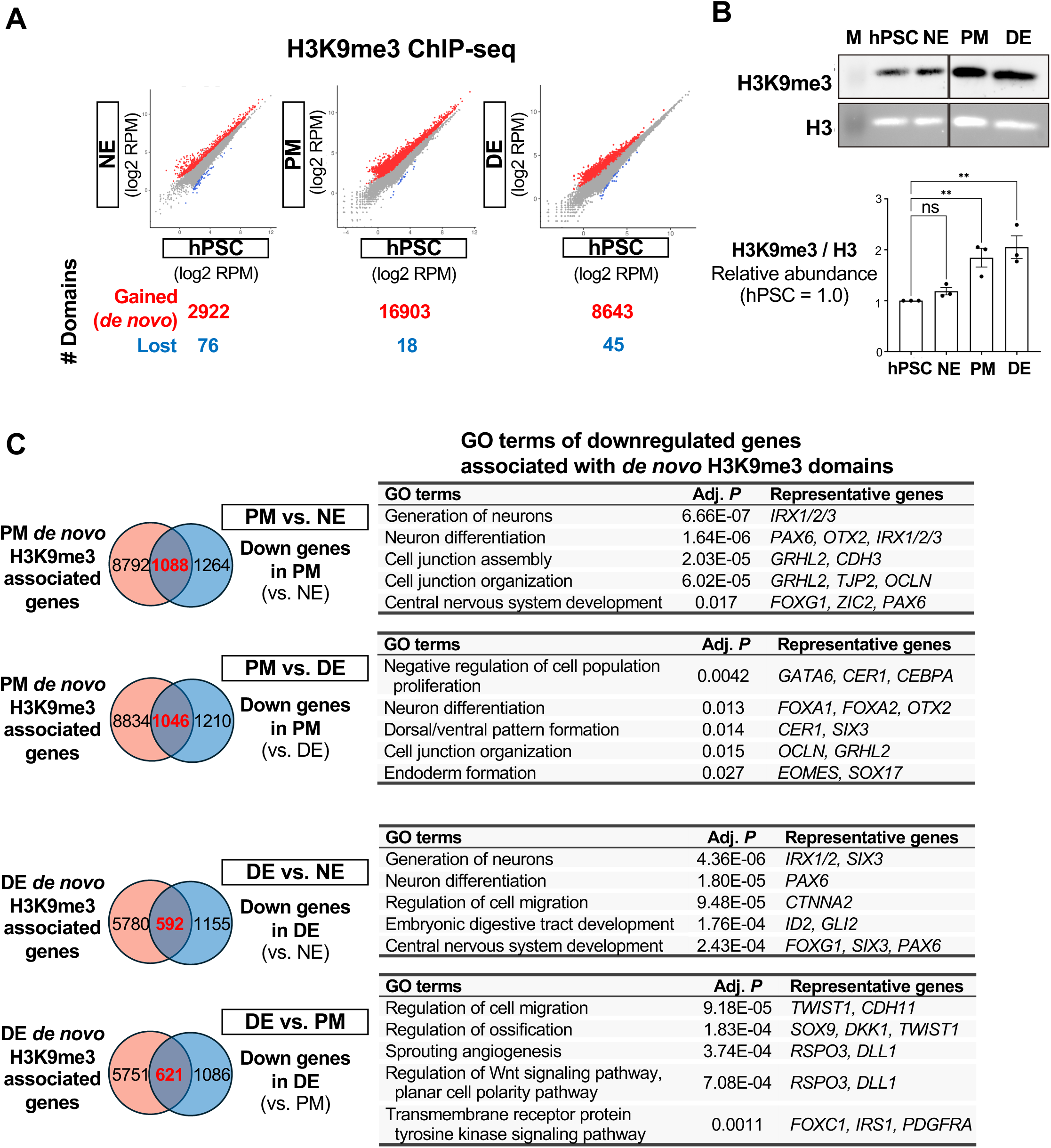
Characterization of H3K9me3 domains and their potential target genes, Related to Figure 1. **(A)** Differential H3K9me3 domain analysis comparing hPSCs and germ layers (n = 2 replicates from 2 independent differentiations; *P* < 0.01 by DESeq2). **(B)** Western blot quantification of relative H3K9me3 signal intensity, normalized to total H3 (n = 3 replicates from 3 independent differentiations, means ± SEM, ** *P* < 0.02, ns = not significant, One-way ANOVA and Dunnett’s test). **(C)** Intersection between genes associated with gained, *de novo* H3K9me3 domains and genes downregulated in each lineage compared to another lineage. GO terms associated with these genes are shown, along with representative genes. Adjusted *P*-values were calculated using Enrichr.

**Figure S3.**
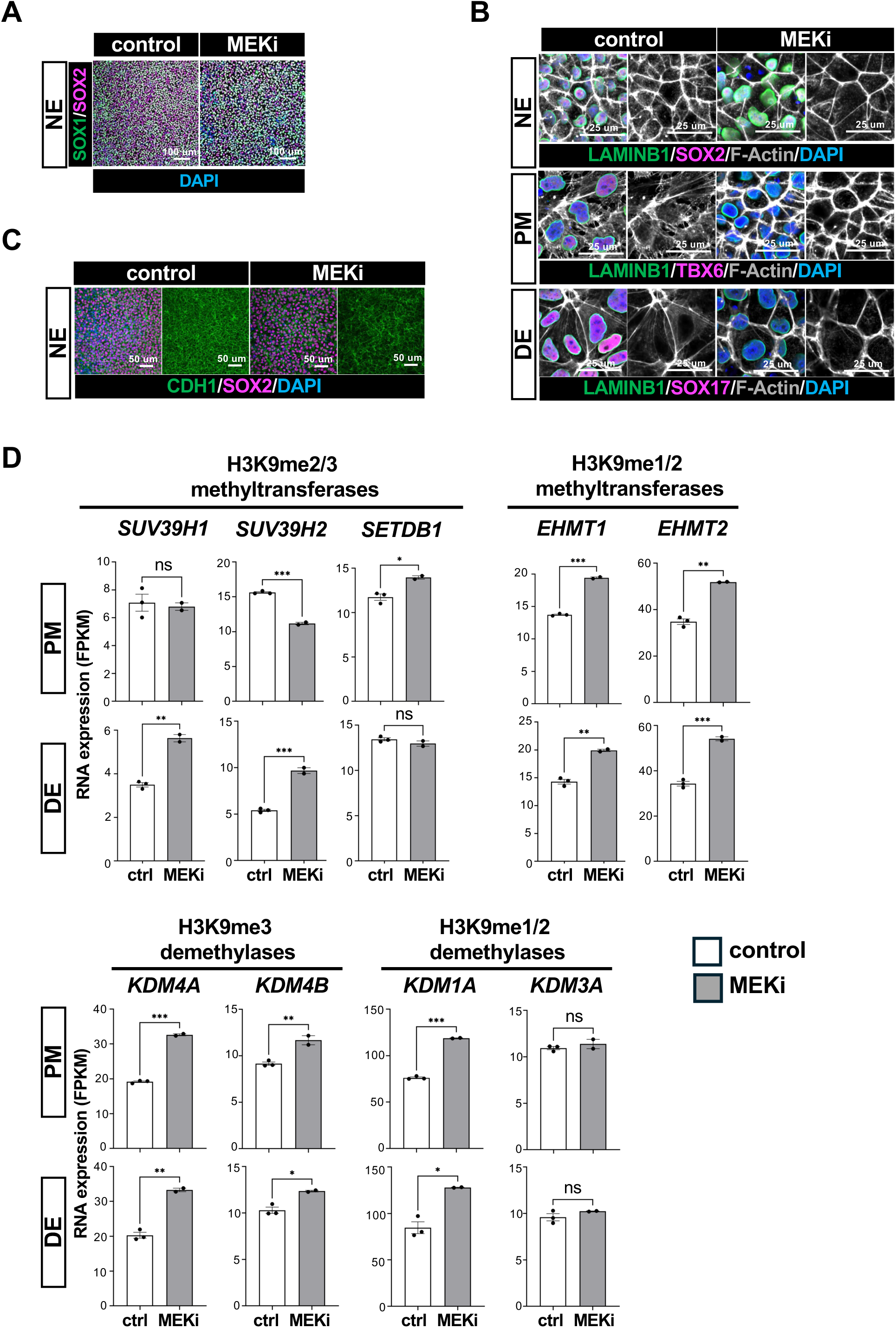
Characterization of MEKi-treated cells, Related to Figure 2. **(A)** Immunostaining showing expression of NE markers (SOX1 and SOX2) in control and MEKi-treated NE. Scale Bars,100 μm. **(B)** Imaging showing the cytoskeleton architecture (F-actin), nuclear shape (LAMINB1), and lineage marker expression (SOX2 for NE, TBX6 for PM, SOX17 for DE) in control and MEKi-treated NE, PM, and DE. Scale Bars, 25 μm. **(C)** Immunostaining showing expression of the CDH1 and SOX2 in control and MEKi-treated NE. Scale Bars, 50 μm. **(D)** Expression of H3K9 methyltransferases and demethylases in control and MEKi-treated PM and DE (n = 2 RNA-seq replicates for MEK-treated cells, n = 3 RNA-seq replicates for control cells; means ± SEM; ** *P* < 0.05; ** *P* < 0.02; *** *P* < 0.001; ns = not significant, Student *t*-test).

**Figure S4.**
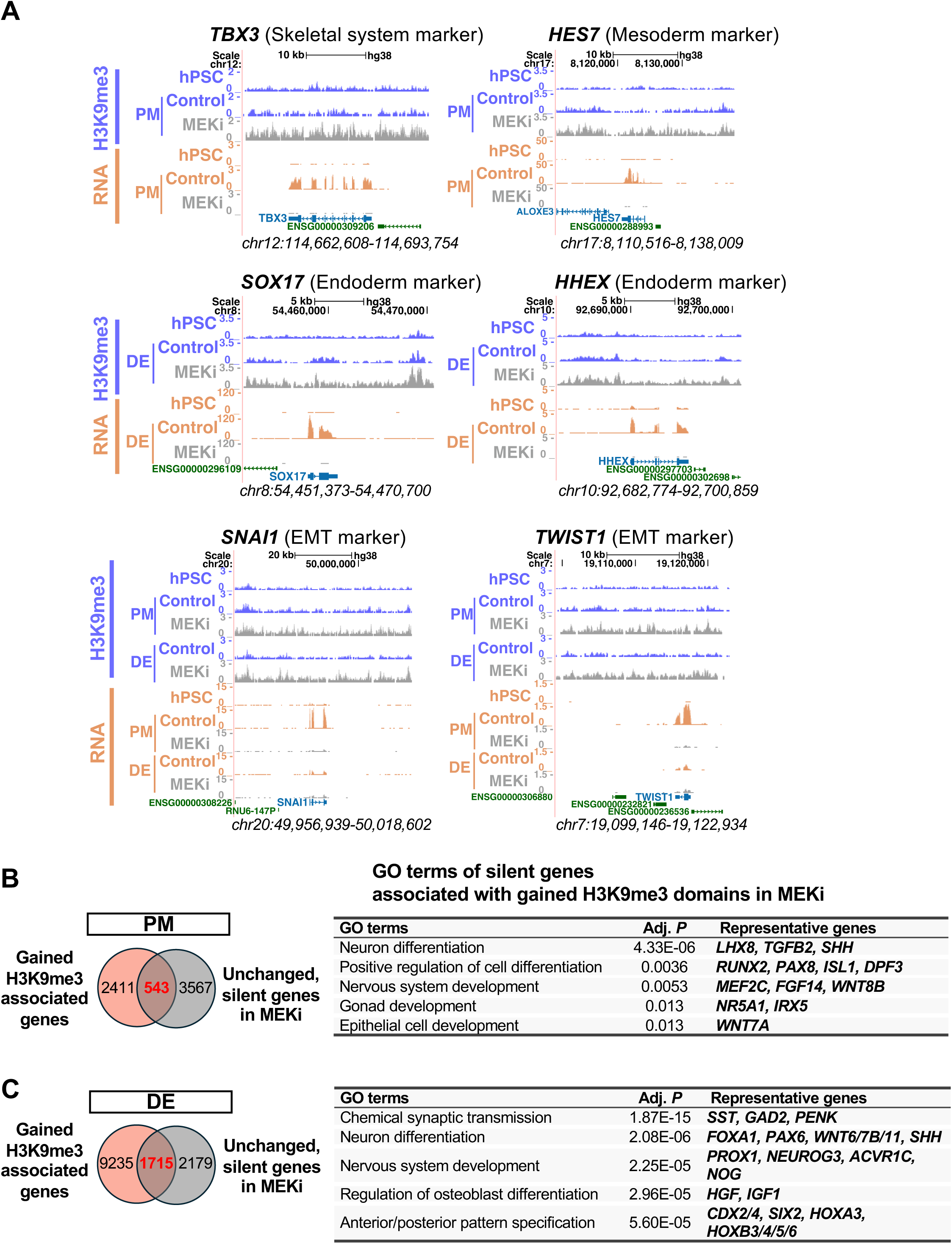
MEK/ERK signaling prevents the establishment of H3K9me3 domains necessary for mesoderm and endoderm differentiation and functions, Related to Figure 4. **(A)** Genome browser tracks displaying H3K9me3 ChIP-seq and RNA-seq data in PM and DE at representative loci of PM, DE, and EMT-related genes. **(B)** Intersection between genes associated with gained H3K9me3 domains and genes unchanged and silent in MEKi-treated PM and DE. GO terms associated with these unchanged and silent genes are shown, along with representative genes. Adjusted *P*-values were calculated using Enrichr.

